# Dual host-bacterial gene expression to study pathogenesis and the regulation of virulence factors in tissue during respiratory infections

**DOI:** 10.1101/2024.07.24.604897

**Authors:** Federico Di Marco, Francesca Nicola, Francesca Giannese, Fabio Saliu, Giovanni Tonon, Stefano de Pretis, Daniela M. Cirillo, Nicola I. Lorè

**Affiliations:** Emerging bacterial pathogens unit, Division of Immunology, Transplantation and Infectious, IRCCS Ospedale San Raffaele, Milan, Italy; Department of Informatics, Systems and Communication, Università degli Studi di Milano-Bicocca, Milan, Italy; Università Vita-Salute San Raffaele, Milan, Italy; Center for Omics Sciences, IRCCS Ospedale San Raffaele, Milano, Italy

**Author notes:** Corresponding author Correspondence to Nicola I. Lorè, Ph.D., Emerging bacterial pathogens unit, Division of Immunology, Transplantation and Infectious, IRCCS Ospedale San Raffaele. Co-Author (Federico Di Marco, Francesca Nicola). Co-Author (Stefano de Pretis, Daniela M. Cirillo, Nicola I. Lorè).

## Abstract

Co-localization of spatial transcriptome information of host and pathogen can revolutionize our understanding of microbial pathogenesis. Here, we aimed to demonstrate that customized bacterial probes can be successfully used to identify host-pathogen interactions in formalin-fixed-paraffin-embedded (FFPE) tissues by probe-based spatial transcriptomics technology. We analyzed the spatial gene expression of bacterial transcripts with the host transcriptomic profile in murine lung tissue chronically infected with *Mycobacterium abscessus* embedded in agar beads. Customized mycobacterial probes were designed for the constitutively expressed *rpoB* gene (an RNA polymerase β subunit) and the virulence factor precursor *lsr2*, modulated by oxidative stress. We found a correlation between the *rpoB* expression, bacterial abundance in the airways, and an increased expression of *lsr2* virulence factor in lung tissue with high oxidative stress. Overall, we demonstrate the potential of dual bacterial and host gene expression assay in FFPE tissues, paving the way for the simultaneous detection of host and bacterial transcriptomes in pathological tissues.

## Introduction

High-resolution technologies are improving our understanding of the complex dynamics of infection processes ^1, 2, 3, 4, 5^. Current approaches lack the simultaneous co-localization of spatial biological information between the host environment and the expression of bacterial virulence factors. Moreover, little information is available on the pathogen’s gene expression and spatial location during infection ^6, 7, 8, 9, 10, 11, 12^. Few recent studies have shown the spatial localization of bacteria in tissues in the context of host transcriptome profiles^9, 10^. Recent progresses have been made in the study of host-virus interaction ^11, 1^, little is known on the expression of specific bacterial virulence genes during infection in pathological tissues. Recent studies related to the spatial localization of bacteria in tissues were limited to the quantification of microbial taxa without providing information on the gene expression profile in individual microbial cells in relation to host transcriptome profiles ^9, 10^. A recent approach based on spatial meta-transcriptomics resolves host-bacteria-fungi interactomes, although this approach lacks the detection of bacterial transcriptomic profiles ^12^. In addition, the potential of an imaging-based approach, i.e. highly multiplexed spatial transcriptomics, was recently shown. The authors profiled cultured bacteria in a wide range of biological and spatial contexts, highlighting the heterogeneity of their gene expression profiles ^13^.

So far, several technical limitations, such as poor permeabilization of the bacterial membrane, lack of polyadenylation of bacterial messenger RNA, and low abundance and stability of bacterial messenger RNA encoding virulence, have hampered the study of bacterial gene expression in tissues using omics technologies ^6, 7, 8, 10, 14, 15^. Current spatial transcriptomic technologies have the potential to overcome these issues by exploiting probe-based strategies to improve the detection of specific RNA transcripts (e.g., those with low abundance or low stability)^16^. In addition, probe-based methods are compatible with formalin-fixed paraffin-embedded (FFPE) tissue blocks, allowing easy access to infected material and to large amounts of samples already deposited in biobanks ^16^.

Here, we hypothesize that existing technological approaches could be adapted to identify the spatial distribution of bacteria and detect virulence factors affected by interactions with inflammatory tissue environments, and vice versa. Among bacterial pathogens, *Mycobacterium abscessus* (*M. abscessus*) is an emerging ubiquitous multidrug-resistant microorganism^17, 18^ that can cause a variety of clinical syndromes/diseases, including the clinical manifestation of non-tuberculous mycobacteria (NTM) lung disease^19^. In NTM disease, it is often unclear whether the pathological outcome in the lung is mediated by the response to *M. abscessus* or by the altered inflammatory microenvironment itself.

Here, to investigate simultaneously the spatially-resolved host and bacterial transcriptional profiles, we employed the probe-based assay (Visium Spatial Gene Expression assay) on FFPE tissue blocks at a resolution of 55 μm (Figure 1A and 1B). We were interested in mycobacterial lung infection, with particular attention to *M. abscessus* ^*17*^; indeed, we utilized tissue blocks from animals chronically infected with *M. abscessus* embedded in agar beads ^20, 21^, as a disease model. To prove the dual spatial gene expression of bacteria and host in tissues, we designed custom probes focusing on two specific mycobacterial transcripts: a constitutive expressed gene, named *rpoB*, that encode for the RNA polymerase β subunit ^22^, and a well-known oxidative stress-modulated bacterial virulence factor precursor, called *lsr2*, which is responsible for encoding a nucleoid-associated protein (NAP)^23 24^.

**Figure 1.**
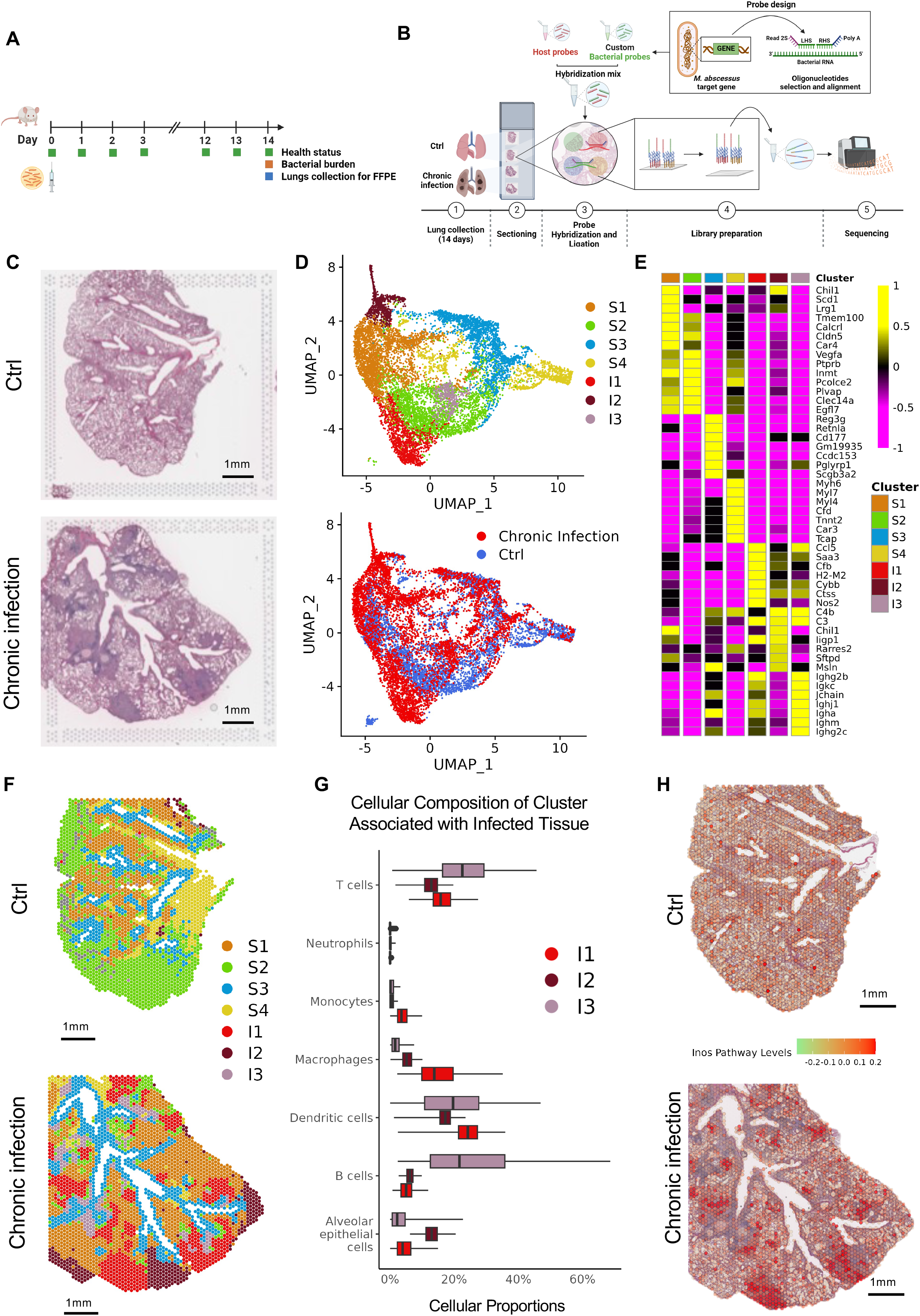
Transcriptional and spatial analysis of not-infected (Ctrl) and chronically infected lungs (Chronic infection). A) Schematic representation of a mouse model of infection (created with BioRender.com) B) Schematic representation of the spatial transcriptomic experiment. (created with BioRender.com) C) H&E staining of Ctrl and chronically infected lungs used for spatial transcriptomic (Scale bar, 1 mm). D) UMAP dimensionality plot of sequenced spots from Ctrl (n=2) and chronically (n=2) infected slices, colour-coded by transcriptional clusters (top) and case (bottom). E) Heatmap showing the expression levels of the top seven transcriptional markers for each cluster, as identified by differential expression analysis. The colour scale represents the average log2 fold change when comparing the cluster to the others. F) Spatial dimensionality plot of Ctrl and Chronic infection slices, colour-coded by transcriptional clusters as in panel D. The spots are arranged according to their spatial coordinates in the tissue sections (Scale bar, 1 mm). G) Boxplot showing the cellular composition distribution of inflammatory clusters (I1, I2 and I3) across Chronic (n=2) Infected slices. H) Spatial plot showing the expression levels of the Inos pathway module in Ctrl and Chronic infection slices (Scale bar, 1 mm).

Using this innovative approach, we found that bacterial gene expression can vary in tissues in different inflammatory areas of the host. The results showed that both *rpoB* and *lsr2* transcripts were detectable and MABS+ areas (spots positive for the presence of *M. abscessus* transcripts) were associated with host inflammatory profiles, including immunoglobulin genes, inflammatory cytokines, macrophagic regulation, and antimicrobial response. Furthermore, *rpoB* expression was higher compared to *lsr2* gene expression and *lsr2* expression was associated with higher levels of the inducible-Nitric Oxide Synthase (iNOS) pathway, responsible for the host response to oxidative stress. The study demonstrates the effective use of a dual bacterial and host gene expression assay in FFPE tissues. Overall, our approach paves the way for the study of microbial pathogenesis in various bacterial diseases by simultaneously detecting the host and bacterial transcriptome in pathological tissues.

## Results and Discussion

### Spatial host gene expression reveals different inflammatory states in lung tissue with chronic *M. abscessus* infection

We used a newly designed dual probe-based method for spatial host-pathogen gene expression (see material and methods) and exploited FFPE tissues from lung of mice displaying chronic infection by *M. abscessus* (ATCC 19977 strain)^17^. Chronic *in vivo* infection with *M. abscessus* was performed using an agar beads murine model. This approach can replicate bacterial persistence in aggregates and planktonic bacilli, which can potentially express diverse bacterial virulence factors as well as a wide range of inflammatory processes in the airways and parenchymal areas ^20, 25, 26^. Mice were infected with an intratracheal injection of 10^5^ colony-forming units (CFU) embedded in agar beads, allowing persistence in the lung of C57BL/6N mice with a sustained bacterial load (Figure 1A). At 14 days post-challenge, these mice display a median bacterial load of around 4 × 10^6^ CFU with a 100% incidence of chronic infection. In this mouse model, *M. abscessus* can persist from 7 to 90 days and is associated with granuloma-like structure areas in the lung, as previously published ^27^.

Therefore, to set up our dual spatial probe-based sequencing, we decided to consider FFPE tissues from one infected mouse (n=2 slices) and one uninfected mouse (n=2 slices) as negative control (Figure 1A and 1B). The two consecutive infected lung slices revealed the histological profile associated with *M. abscessus* persistence, including inflammatory foci characterized by infiltrating macrophages or early granuloma-like structures (Figure 1C, Supplementary Figures 1 and 4).

The transcriptome profiles of the infected lung exhibited distinct expression patterns compared to the control mouse, as shown in the unsupervised cluster analysis (Figure 1D, 1E, 1F, Supplementary Figures 1, 2, and 3). Moreover, we identified a total of four clusters shared between control and infected mice (named S1, S2, S3, and S4), and three clusters located in the chronically Infected mouse (named I1, I2, and I3) (Figures 1 and 1E, Supplementary Figures 1, 2 and 3). Clusters located in chronically infected mice colocalized with pathological areas, including within (Cluster I1) or surrounding (Cluster I2) tissue with infiltrating macrophages/ granuloma-like structures, or with submucosal inflammatory cells (Cluster I3), as observed in terms of H&E staining (Figure 1F, Supplementary Figures 1 and 4). Similar findings in terms of spatial distribution were confirmed by our dual spatial probe-based assay when different and consecutive FFPE tissue sections were analyzed for both groups (Supplementary Figures 1, 2, 3). We carried out a deconvolution approach to define the proportion of cell types for each spot in the four slices using the Robust Cell-Type Decomposition (RCTD) method ^4, 28^, and an annotated single-cell RNAseq reference dataset from Xu et al. ^4^ (GEO: GSE190225) comprehending 12 cell types from Mouse lung infected with *Klebsiella pneumoniae*. The deconvolution results were shown as proportions of 12 cell types in each cluster (Supplementary Figure 5). Shared clusters (S1, S2, S3, and S4) were mainly composed of stromal cells, such as alveolar, club, endothelial, and fibroblast cells. In contrast, the three clusters within the chronically Infected mouse (I1, I2, and I3) displayed higher proportions of inflammatory cells, including macrophage, monocyte, B, and T cells (Supplementary Figure 5). Those cellular profiles were in line with Gene Ontology terms associated with cluster markers (Supplementary Figures 5, 6 and Supplementary data).

Following this, we focused our attention on cluster I1, which presented a high-level proportion of granuloma-associated cells, such as macrophages, dendritic cells, club cells, and T cells ^5, 29, 30^ (Figure 1G). To further confirm the host spatial molecular characterization of the I1 cluster, we determined the spatial distribution of gene expression markers (selected from Gene Ontology terms) associated with host response to *M. abscessus*, such as the iNOS or reactive oxygen species (ROS) pathways. We confirmed that the signal level of both iNOS and ROS pathways were upregulated in the infected mouse compared with the control group (Supplementary Figures 7 and 8) and co-localized with the granuloma-like structure, as shown in Figure 1H and Supplementary Figures 4, 5, and 6.

### Spatial bacterial gene expression detects microbial expression patterns in lungs with chronic *M. abscessus* infection

Next, we determined the feasibility of assaying bacterial and host probes in the same tissue section using our dual spatial probe-based sequencing approach. In addition to the 18k host gene targeted by the commercial kits, we designed and developed 3 pairs of custom probes per gene, targeting two distinct bacterial transcripts. We were interested in showing the effectiveness of this approach in mycobacterial infection, with particular attention to *M. abscessus*. Therefore, we focused on two distinct mycobacterial transcripts: one constitutive expressed gene, named *rpoB*^22^, and a well-known precursor of bacterial virulence factor modulated by oxidative stress, referred to as *lsr2*^23,24^. We also refined the Visium protocol by incorporating our new probes into the modified permeabilization protocol to improve probe hybridization due to the mycobacterial membranes (see Material and Methods). The expression of the selected bacterial targets, *rpoB* and *lsr2*, was spatially detected (Figure 2A and Supplementary Figure 9). To validate the detection obtained with our *M. abscessus* probes, we compared *M. abscessus* localization by spatial transcriptomics assay with *M. abscessus* detection assays by orthogonal Kinyoun Stain (Supplementary Figure 9) and ad hoc immunofluorescence staining (Figure 2B) on consecutive tissue sections. We observed that airways and submucosal airway areas displayed a higher number of bacteria (Figure 2A I) compared to parenchymal or granuloma-like structure areas (Figure 2A II, III) of uninfected control regions (Figure 1A IV) and control mouse (Supplementary Figure 10). Overall, these results demonstrate that the selected bacterial probes and the refined protocol can be used to detect bacterial transcripts in infected organs co-localized with host transcriptome profiles.

**Figure 2:**
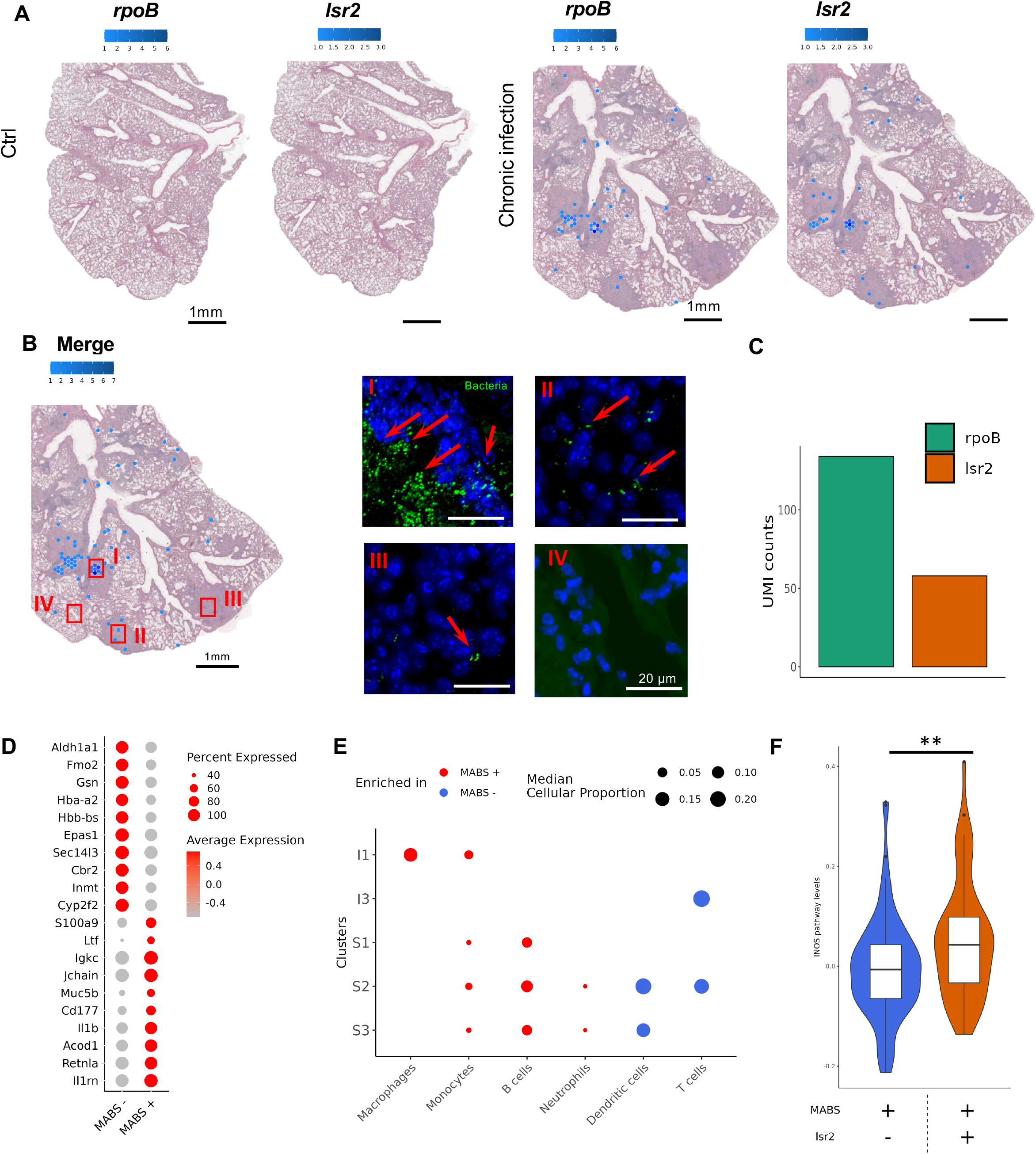
Spatial and quantitative analysis of *rpoB* and *lsr2* probes in Ctrl and Chronic infection slices. A) Spatial plot showing the UMI counts of rpoB and *lsr2* probes in Ctrl and Chronic infection representative slices. The colour scale indicates the number of UMIs per spot. (Scale bar, 1 mm). B) Spatial plot showing the summed UMI counts of *rpoB* and *lsr2* probes in Chronic infection slices (n=2). The colour scale indicates the total number of UMIs per spot (Scale bar, 1 mm). (I-IV) Representative images of immunofluorescence staining in infected mouse for *M. abscessus* (with specific antibody against *M. abscessus*, stained in green, and with Hoechst 33342, Trihydrochloride, stained in blue) in airways and submucosal airways (DI), in granuloma-like structure areas (D II, III), and area of not inflamed parenchyma (D IV) all captured in the successive slice of the infected sample used for spatial transcriptomics (Scale bar, 20 μm). C) Barplot showing the gathered UMI counts of *rpoB* and lsr2 probes in Chronic infection slices. The y-axis shows the number of UMIs. The bars are colour-coded by the respective probe. D) Bubbleplot showing the differentially expressed genes in MABS+ (positive) and MABS- (negative) spots in Chronic infection slices (n=2). The color scale shows the log fold change of the genes. The size of the bubbles indicates the percentage of spots expressing a gene. E) The bubble plot illustrates the significantly different cellular populations between MABS+ (positive) and MABS- (negative) spots, categorized by clusters. Red dots indicate enrichment in MABS-positive spots, while blue dots indicate enrichment in MABS-negative spots. The size of each dot corresponds to the median cellular proportion observed for the specific cluster. F) Violin plot for the levels of INOS pathway grouped according to MABS+lsr2- or MABS+lsr2+ Infection (generated with chronic infection slices, n=2) (Mann-Whitney test,⍰**p-value⍰<0,01).

### Dual spatial host bacterial gene expression reveals *M. abscessus* and its virulence factor *lsr2* are associated with different inflammatory states in lung chronic infection

Next, we investigated the distribution of *M. abscessus* transcripts profile in the two consecutive tissue sections from the infected mouse. In these sections, 2.15 % of spots had an *M. abscessus* transcriptional signal (MABS+ areas, with positive spots for the presence of either *rpoB* or *lsr2* or both transcripts), with highly reproducible capture of *M. abscessus* gene spots (67 and 63 bacterial spots out of 3176 and 2875 total spots in slice C1 and D1, respectively) (Supplementary Figure 9A). In addition, we found that UMI counts were higher for the constitutively expressed *rpoB* gene than for the *lsr2* gene, and the overall numbers were similar between two different sections (Figure 2C and Supplementary Figure 11A).

Our approach allowed us to perform dual spatial host and *M. abscessus* gene expression analysis to identify changes in host gene expression induced by bacterial presence in lung cells at a resolution of 55 μm. We compared host gene expression patterns between the *M. abscessus* positive (MABS+, positive spots for the presence of *M. abscessus* transcripts) and negative (MABS−, negative spots for *M. abscessus* transcripts) spots. In the top 10 up- and down-regulated genes, we found upregulation of a specific immunoglobulin gene (*Igkc*), inflammatory cytokines (*Il1b* and *Il1rn*), macrophagic regulation (*Retnla* and *Acod1*) as well as neutrophilic (*Cd177*) and antimicrobial response (*S100a9*) in the MABS+ spots (Figure 2D) (differential expression analysis used the Mann-Whitney test, p-value⍰<⍰0.05). In addition, we compared cellular composition associated with MABS+ and MABS-spots among clusters (Figure 2E, see Methods). MABS+ spots were significantly enriched in Macrophages and Monocytes within Cluster I1, colocalized with granuloma-like structures, while Monocytes, B cells, and Neutrophils were mainly enriched in MABS+ spots within Cluster S1, S2 and S3. Only Dendritic and T cell proportions were mainly present in MABS-spots within I3, S2, and S3 clusters (Figure 2E).

Moreover, we hypothesized that bacterial probes targeting either a constitutively expressed gene, *rpoB*^22^, or a bacterial virulence factor precursor, *lsr2*, may differently be expressed in diverse host inflammatory areas. As mentioned, *Isr2* is a virulence factor precursor critical for *M. abscessus* in oxidative stress condition and for macrophage internalization in mycobacterial infection ^23 24 31^. Therefore, we determined whether diverse host tissue immune responses (e.g. lung areas with different oxidative stress states) in the bacterial positive spots (MABS+) could influence the expression of *lsr2* transcripts differently. When we examined spatial gene expression in the infected lung related to oxidative stress processes, such as the nitric oxide biosynthetic pathway, we confirmed that the *lsr2*+ positive spots among MABS+ spots (MABS+ *lsr2*+) were significantly enriched in the expression of iNOS pathways than lsr2-negative spots (MABS+ *lsr2-*) (Figure 2F, adjusted p <0.01 and Supplementary Figure 11B). Overall, we confirmed that lung areas with high oxidative stress conditions were co-localized with the expression of the *lsr2* virulence factor precursor.

## Conclusions

Our results show that bacterial probes are suitable for the spatial localization of bacteria in inflamed and not-inflamed lung tissues. Moreover, our study highlights the power of this method for detecting bacteria and their virulence factors under modulation by the local host environment, such as mycobacterial *lsr2* expression and the inducible nitric oxide host response. In addition, thanks to our approach and data on virulence factor precursor expression of the *lsr2* gene in pathological tissue, we can confirm its relevance both as a potential vaccine candidate against *M. abscessus* infection and as well as a bacterial target required for productive mycobacteriophage infection, as already published^23, 24, 32^. Defining the interaction between *M. abscessus* and the lung regarding the organization and spatial distribution of bacteria and infected niches is needed in treatment decision-making, requiring a deeper understanding of the complex mycobacterial-host interaction in pathological tissues and associated lung diseases.

Our approach deciphers the simultaneous and spatially distributed expression of mycobacterial and host genes in tissues and can decode the spatial pathological complexity of chronic lung disease caused by *M. abscessus* interactions. This study evidence supports that our approach is consistent and can be scaled up to target several bacterial transcripts. Moreover, our data suggest that *M. abscessus* can be associated with specific cell types based on tissue spatial localization, such as the mucosal barrier or within granuloma-like structures. Therefore, increasing the number of biological replicates and scaling up the bacterial probes could allow for a more accurate dissection of cell type/bacteria interactions, including specific host gene expression patterns associated with bacteria. In the future, using pools of probes targeting the overall *M. abscessus* transcriptome we will assess the diverse bacterial population states within distinct geographical areas of the mouse lung, simultaneously with host transcriptomic profiling. In addition, an increased number of biological replicates and a deeper sequencing of the bacterial probes will highlight differences in spatial localization among bacterial transcripts. Our custom approach, based on the design of a unique set of probes to define specific bacterial transcriptome profiles, is not limited to mycobacterial infections. In fact, it can be broadly applied to other infectious agents affecting the lung or other organs, such as *Pseudomonas aeruginosa, Klebsiella pneumoniae, Salmonella enterica typhi*, among others. In conclusion, our results demonstrate the power of the dual host-bacterial gene expression assay for studying microbial pathogenesis in various bacterial diseases and pave the way for future research on how inflammatory environments modulate pathogen virulence and vice versa.

## Acknowledgements

The authors thank Fiocchi A. and Guidotti L. (ANIMAL HISTOPATHOLOGY, IRCCS Ospedale San Raffaele, Milano, Italy) for the mouse histopathology preparations and the San Raffaele Microscopy facility (Alembic, Ospedale San Raffaele, Milano, Italy) for immunofluorescence images acquisition. In addition, the authors are grateful to Morgan GSK (Postdoctoral fellow, IRCCS Ospedale San Raffaele, Milano, Italy) for the critical reading and English grammar correction of the manuscript.

## Author contributions

FD, FN, SdP and NIL conceptualized the study. SdP and NIL supervised the study. FN, FS, NIL performed processing protocols and in vivo experiments. FN, FG, FS, NIL performed visium spatial library sequencing. Data was processed, curated, and visualized by FD and FN under the supervision of SdP and NIL, and was analysed by FD and SdP. The manuscript was drafted by FD, FN, SdP, NIL and was reviewed and edited by FG, GT, CD.

## Competing interests

All other authors declare no competing interests.

## Notes

### Competing Interest Statement

The authors have declared no competing interest.

