## Supplementary Figures for "Dual host-bacterial gene expression to study pathogenesis and the regulation of virulence factors in tissue during respiratory infections"

### SUPPLEMENTARY FIGURE 1

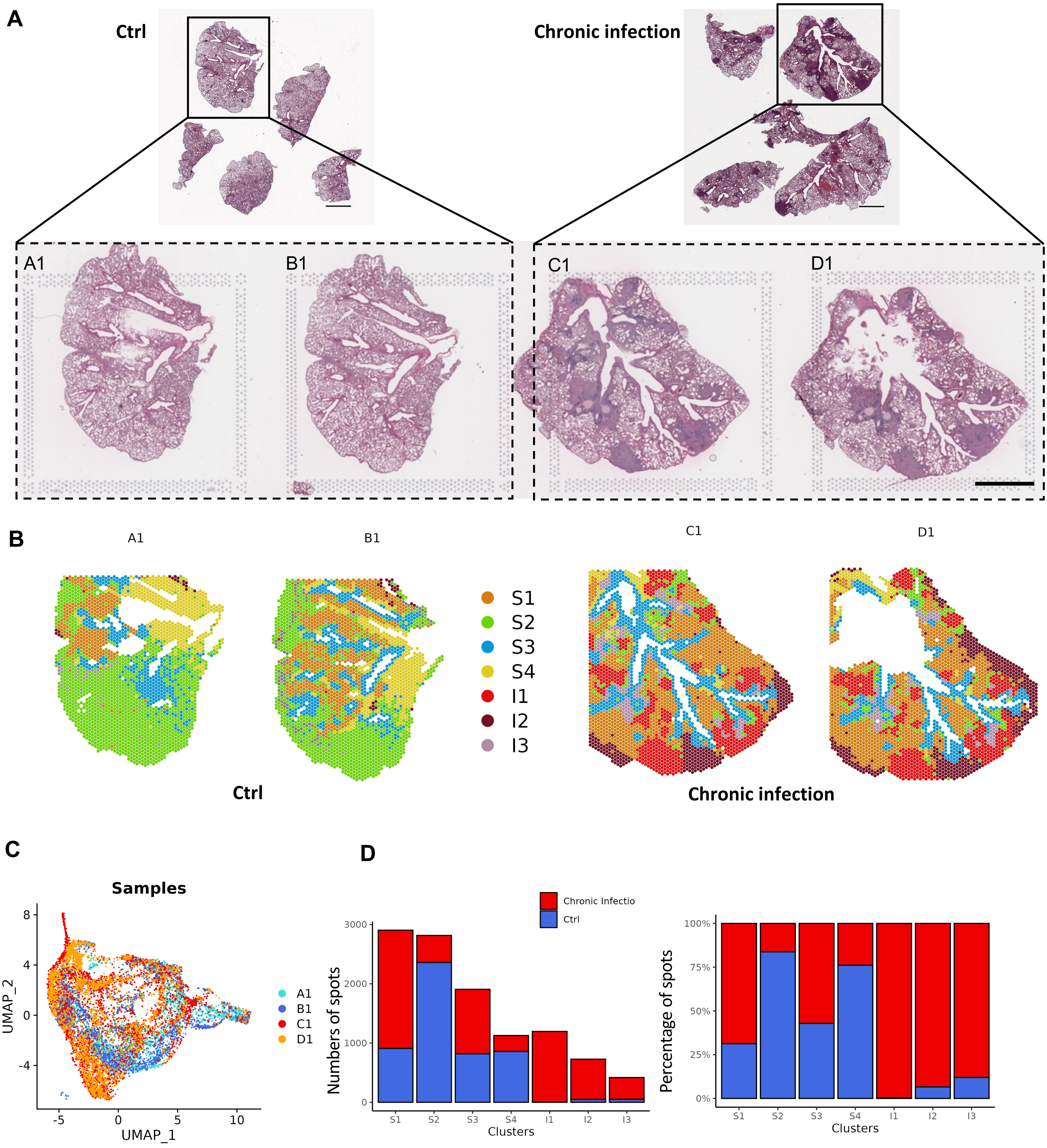


**Supplementary Figure 1: A)** View of 5 lobes in tissue blocks from control and chronically infected mouse stained with H&E. We selected one lobe from each mouse to perform the experiments. At the bottom, there is a summary of four lung tissue slides, each stained with H&E, covering the four capture areas on the Visium Spatial Transcriptomic slide (Scale bar, 2mm). **B)** Spatial dimensionality plot of Ctrl (A1 and B1) and Chronic infection slices (C1 and D1), color-coded by transcriptional clusters as in Figure 1D. The spots are arranged according to their spatial coordinates in the tissue sections. **C)** UMAP dimensionality plots of sequenced spots from A1, B1, C1, and D1. **D)** Barplots showing the number of spots associated to each cluster and the ratio (Percentage) of spot associated to each clusters color coded by case.

### SUPPLEMENTARY FIGURE 2

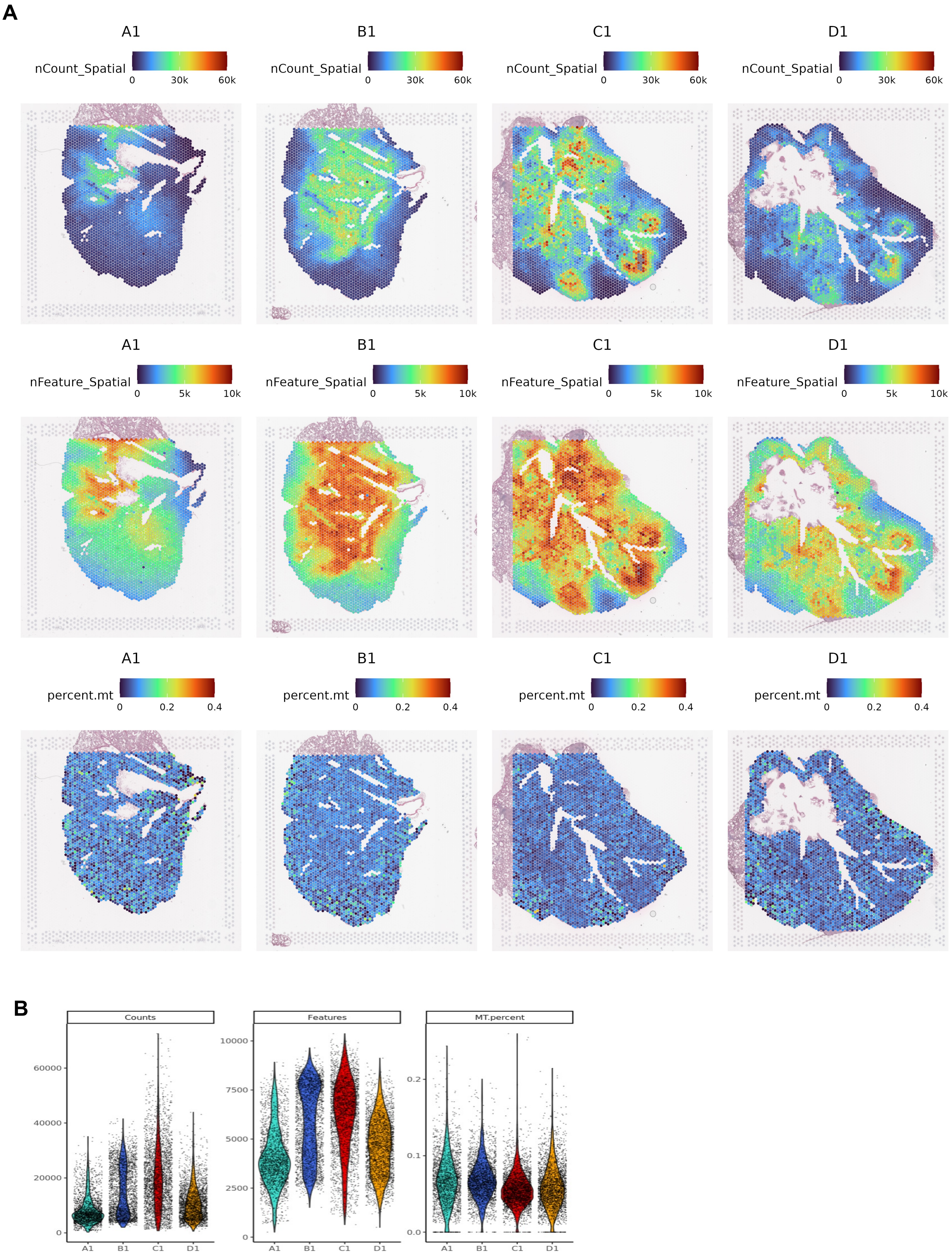


**Supplementary Figure 2: A)** Spatial Plot of Ctrl (A1 and B1) and Chronic Infection (C1 and D1) UMI counts, Feature counts and mithocondrial gene percentages per spot. The spots are arranged according to their spatial coordinates in the tissue sections. **B)** Violin plots of A1, B1, C1, D1 UMI counts, Feature counts and mithocondrial gene percentages per post.

### SUPPLEMENTARY FIGURE 3

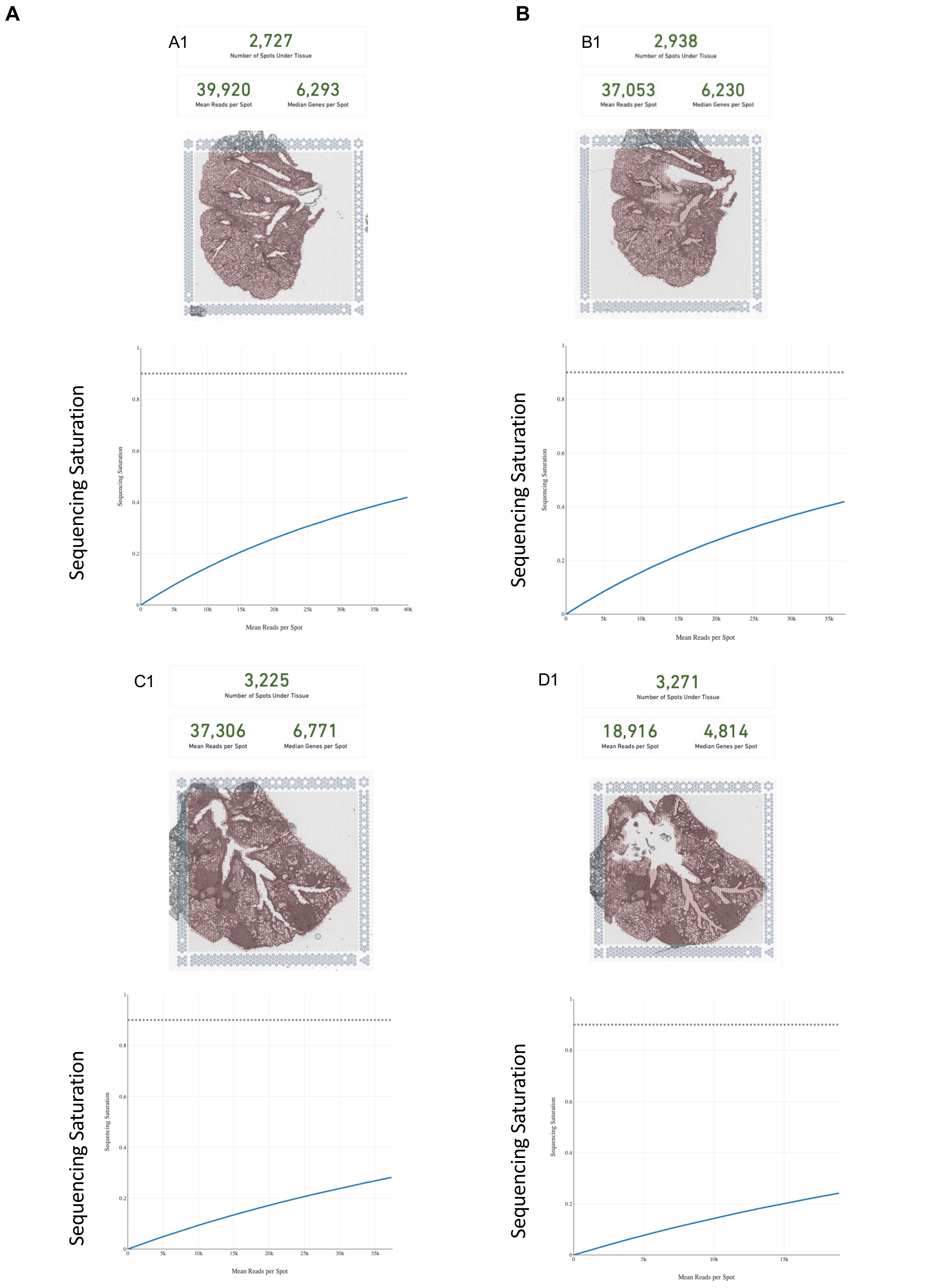


**Supplementary Figure 3:** Space Ranger summary of sequencing (**A**) Batch 1 and (**B**) Batch 2

SUPPLEMENTARY FIGURE 4

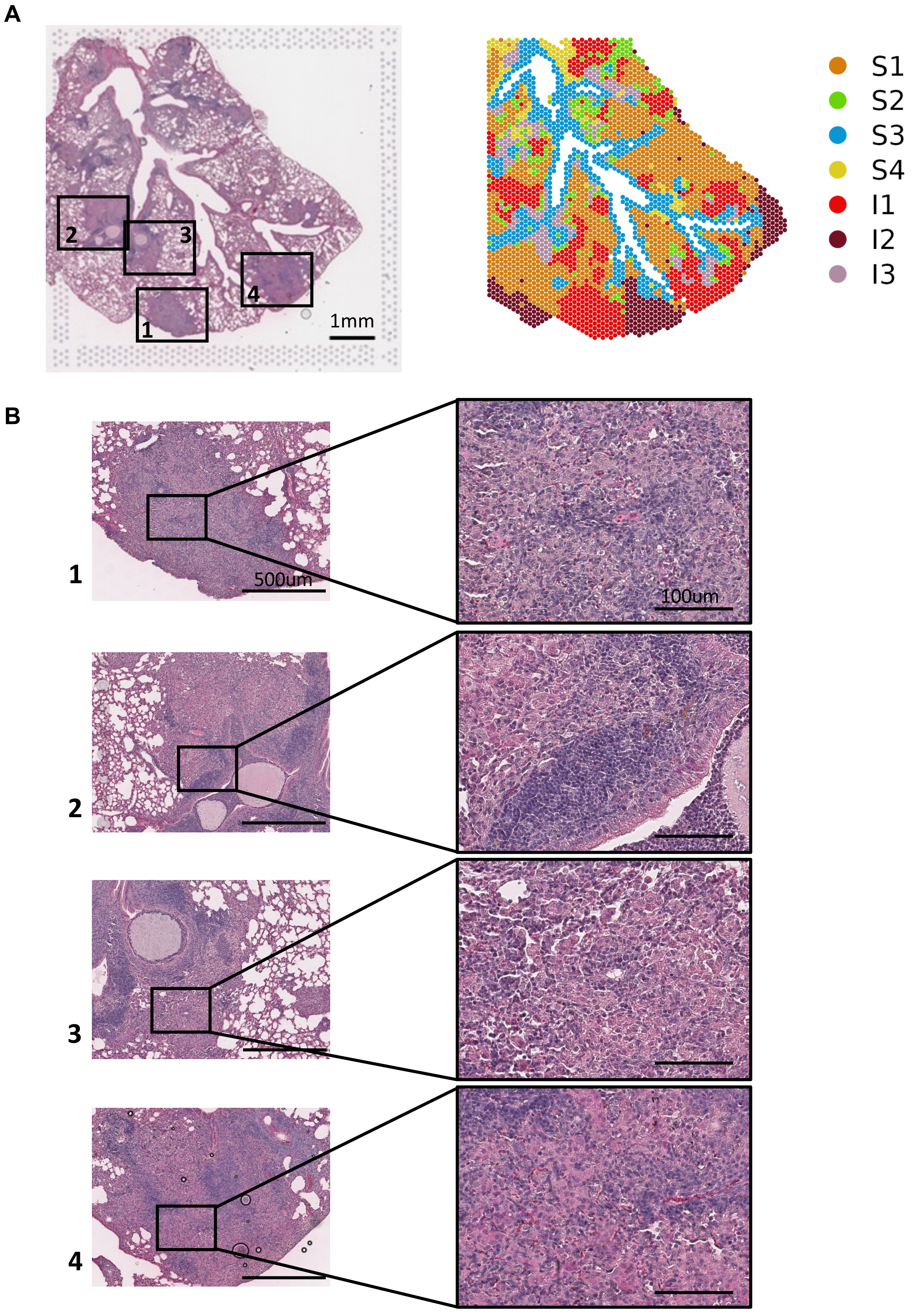


**Supplementary Figure 4: A**) H&E staining and spatial dimensionality plot of Chronic infection slice, color-coded by transcriptional clusters as in Figure 1D (Scale bar, 1 mm). **B**) Representative images of H&E staining, highlighting inflammatory foci characterized by infiltrating macrophage or early granuloma-like structures (1-4) (Scale bar, 500µm and 100µm).

**SUPPLEMENTARY FIGURE 5**

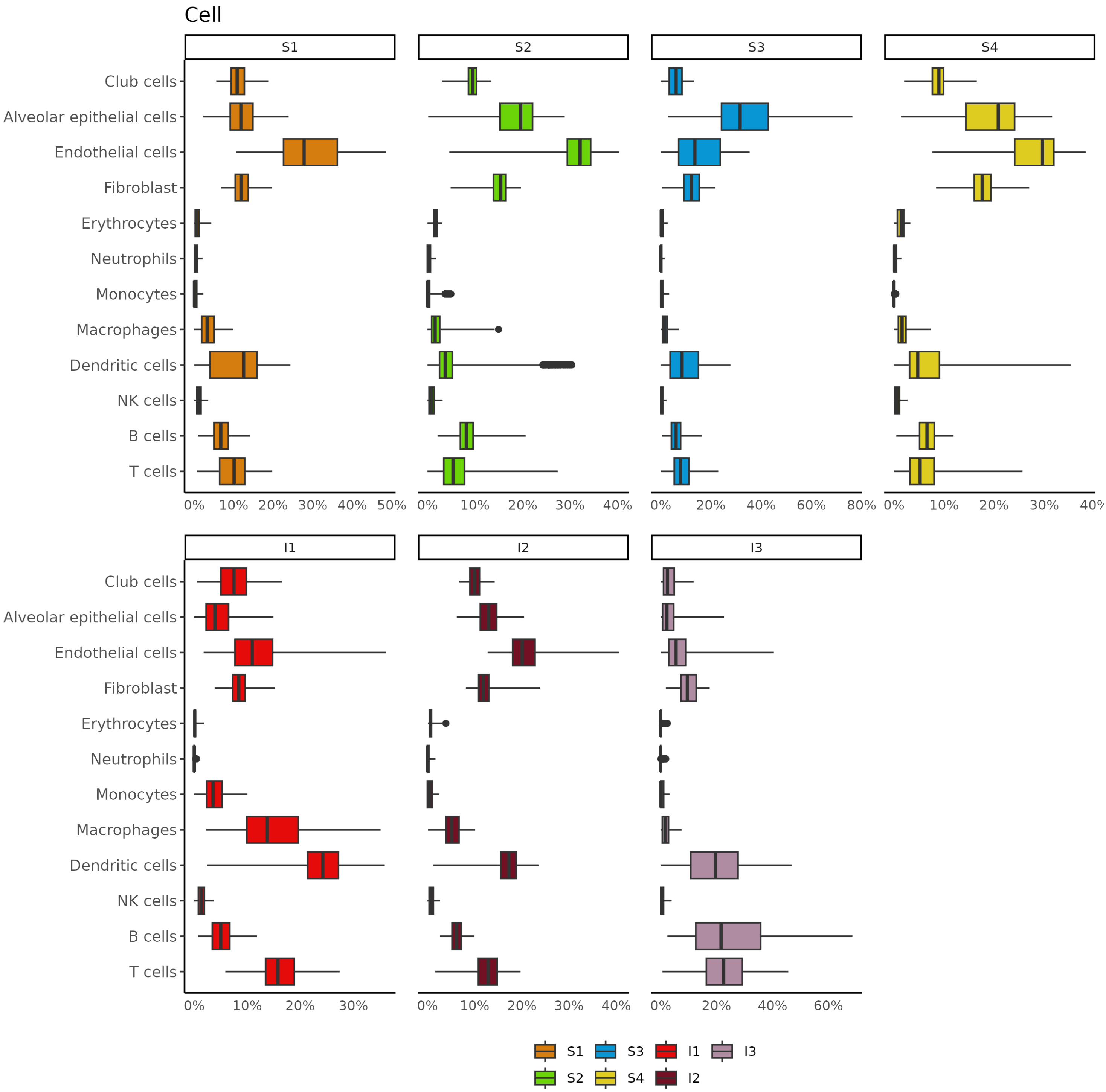


**Supplementary Figure 5: A**) Boxplot showing the cellular composition distribution stratified by clusters.

**SUPPLEMENTARY FIGURE 6**

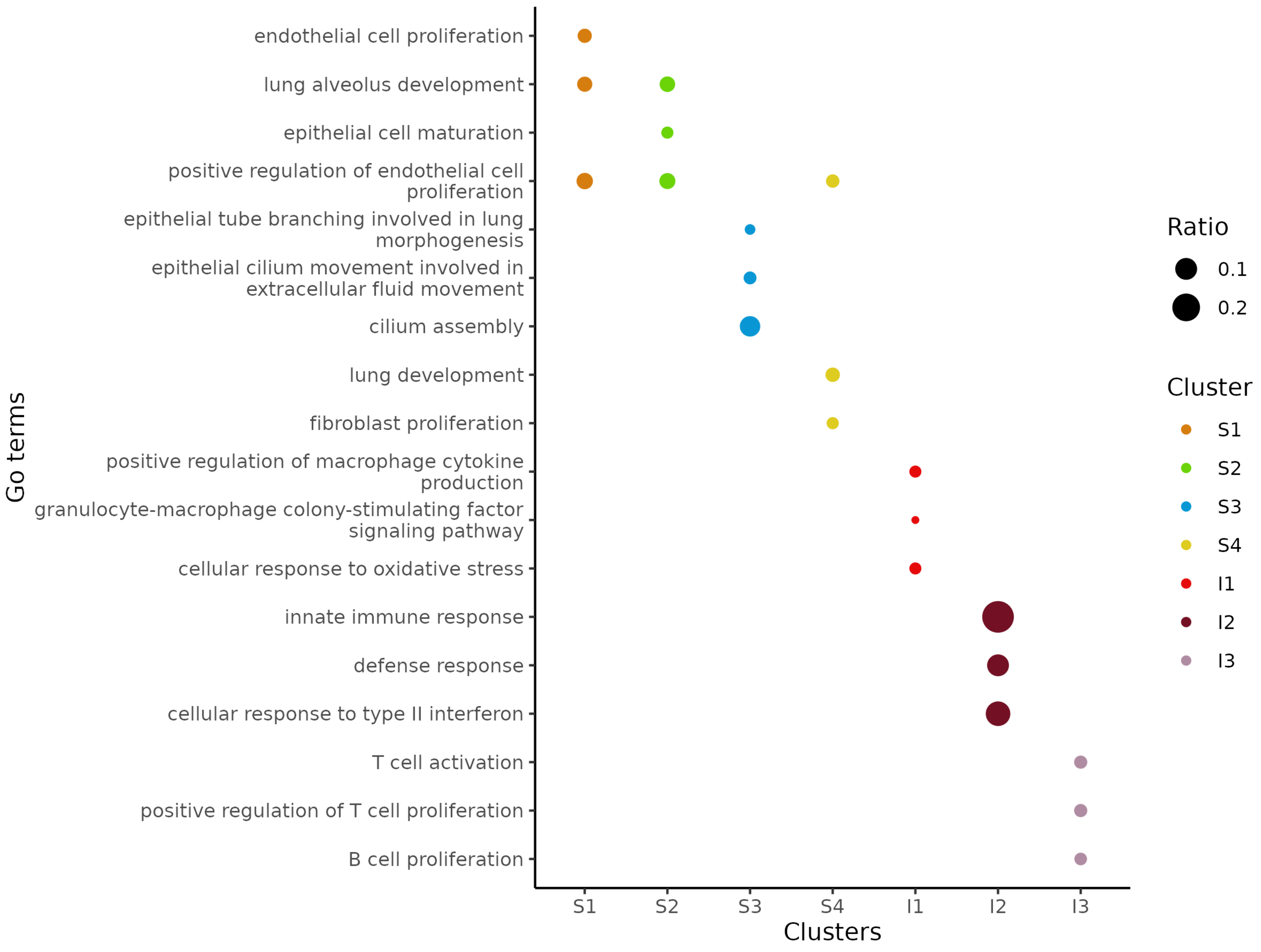


**Supplementary Figure 6: A**) The dot plot displays a selection of enriched Gene Ontology (GO) terms derived from cluster markers. Dots are color-coded according to their respective clusters, and their size represents the ratio of enriched genes associated with each specific GO term.

**SUPPLEMENTARY FIGURE 7**

**A**

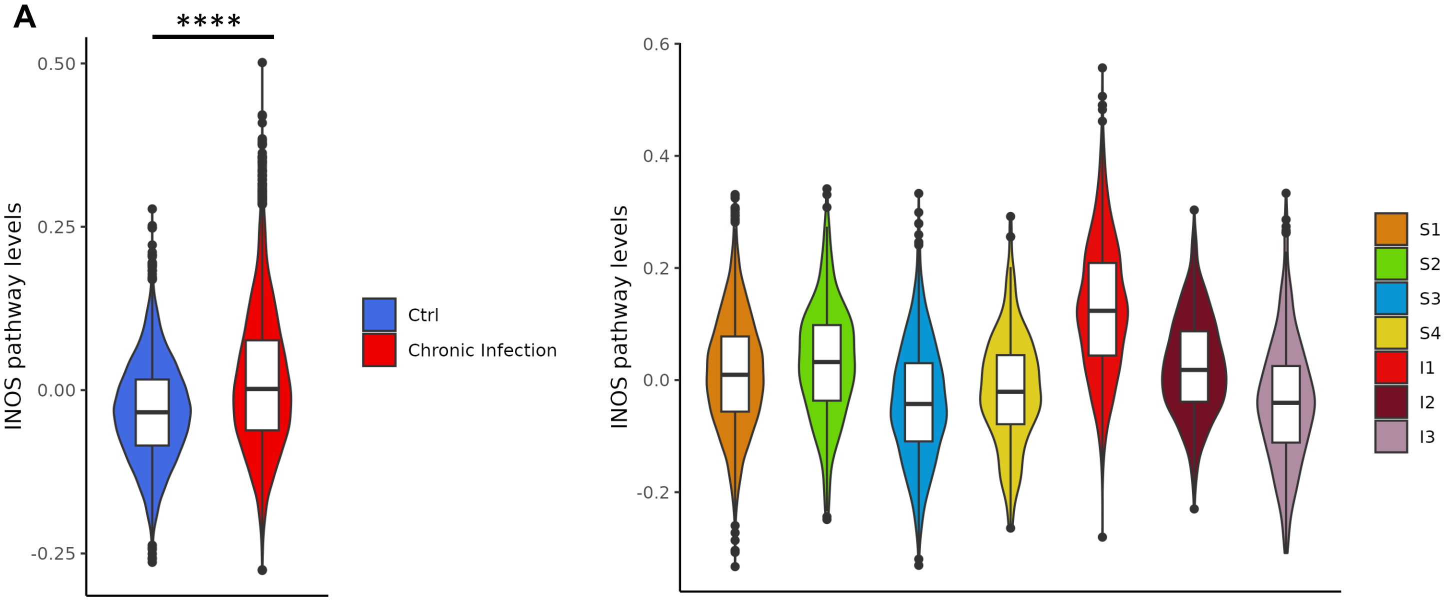


**B**

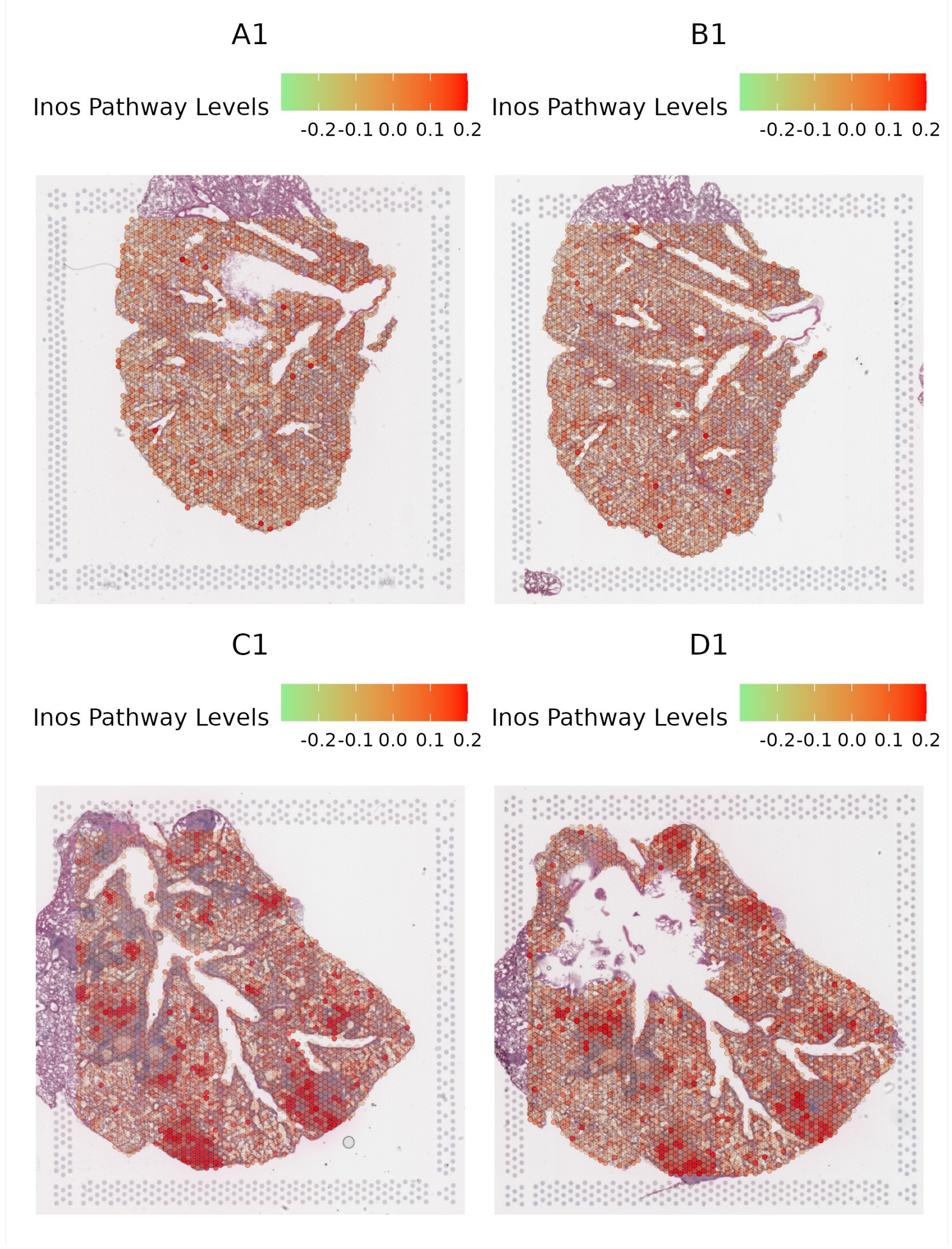


**Supplementary Figure 7: A)** Violin plot for the levels of INOS pathway grouped according to Ctrl vs Infection on the left and according to Clusters on the right (Mann-Whitney test **** P<0,0001). **B)** Spatial plot showing the expression levels of the iNOS pathway module in Ctrl and Chronic infection slices.

**SUPPLEMENTARY FIGURE 8**

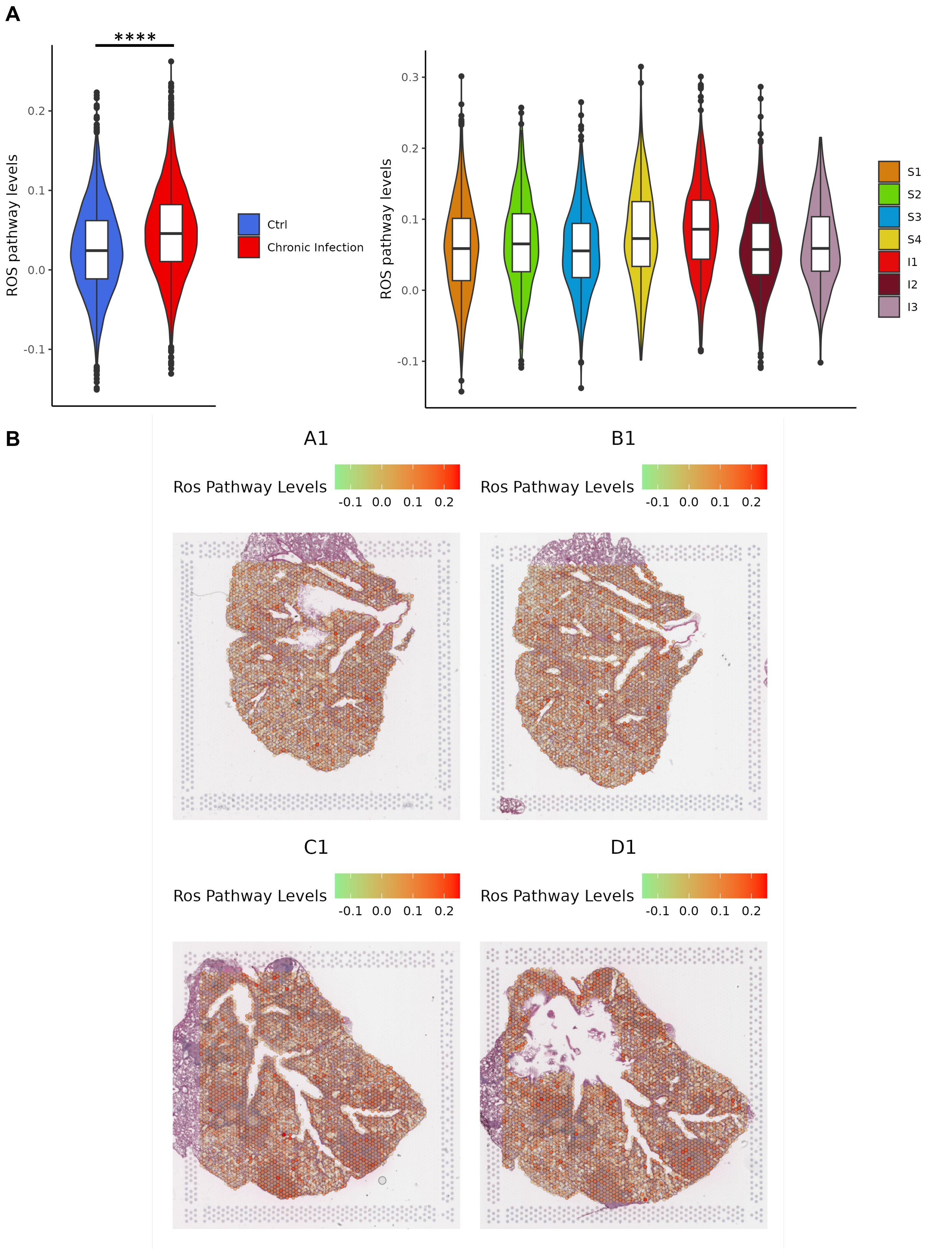


**Supplementary Figure 8: A)** Violin plot for the levels of ROS pathway grouped according to Ctrl vs Infection on the left and according to Clusters on the right (Mann-Whitney test **** P<0,0001) B) Spatial plot showing the expression levels of the ROS pathway module in Ctrl and Chronic infection slices.

### SUPPLEMENTARY FIGURE 9

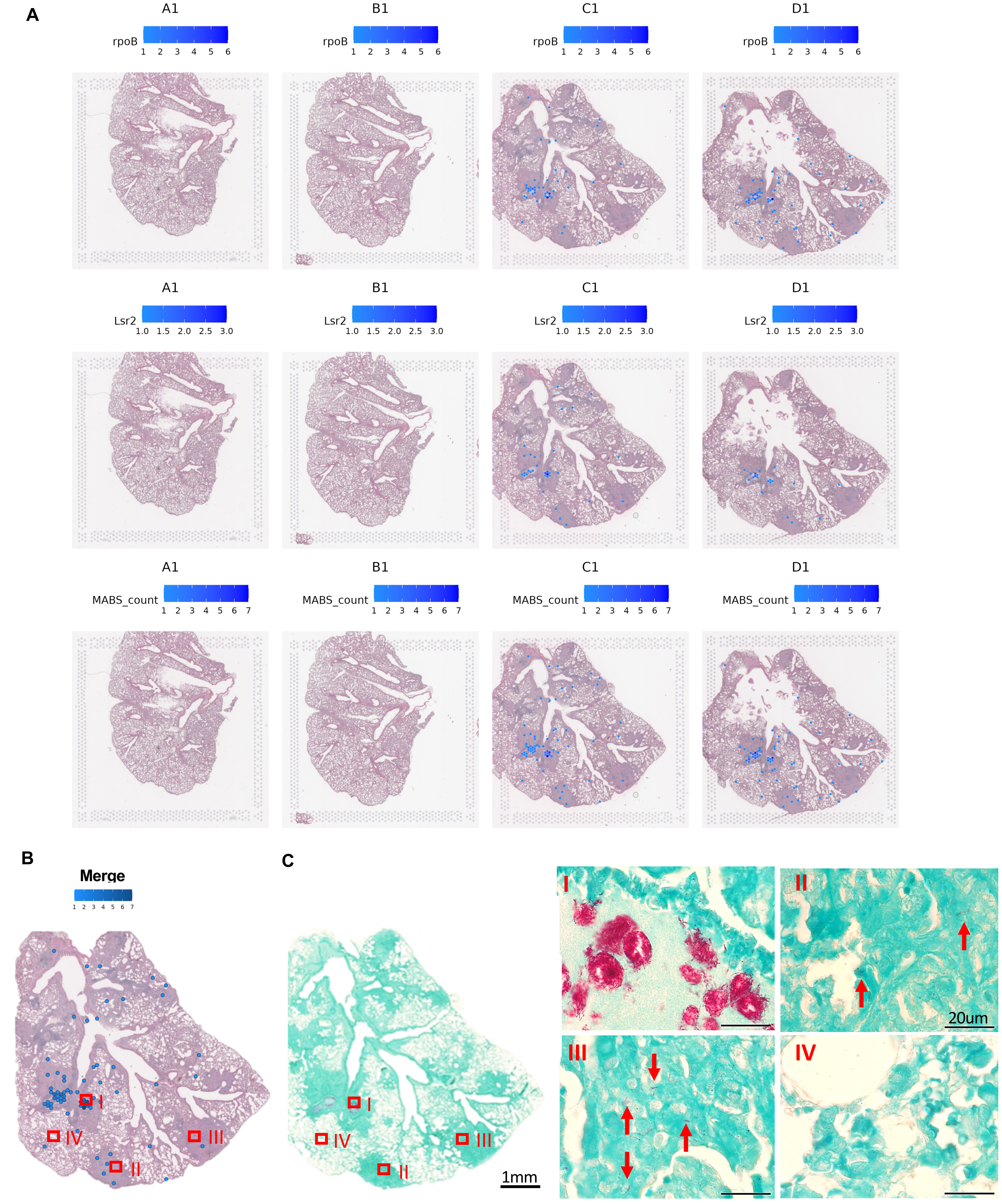


**Supplementary Figure 9:** A) Spatial plot showing the UMI counts of rpoB, lsr2, and summed probes in Ctrl (A1 and B1) and Chronic infection slices (C1 and D1). The color scale indicates the number of UMIs per spot. **B**) Spatial distribution of M. *abscessus* positive spots indicating summed counts for rpoB and lsr2 probes in chronically Infected mouse lung sections. **C**) Representative images of AFB staining kit for histology for M. *abscessus* stained with fuchsin highlighting (100x) MABS distribution in airways and submucosal airways (C I), in granuloma-like structure areas (indicated by arrows) (C II, III), and undetected in the area of not inflamed parenchyma (C IV) (Scale bar, 1 mm and 20μm).

### SUPPLEMENTARY FIGURE 10

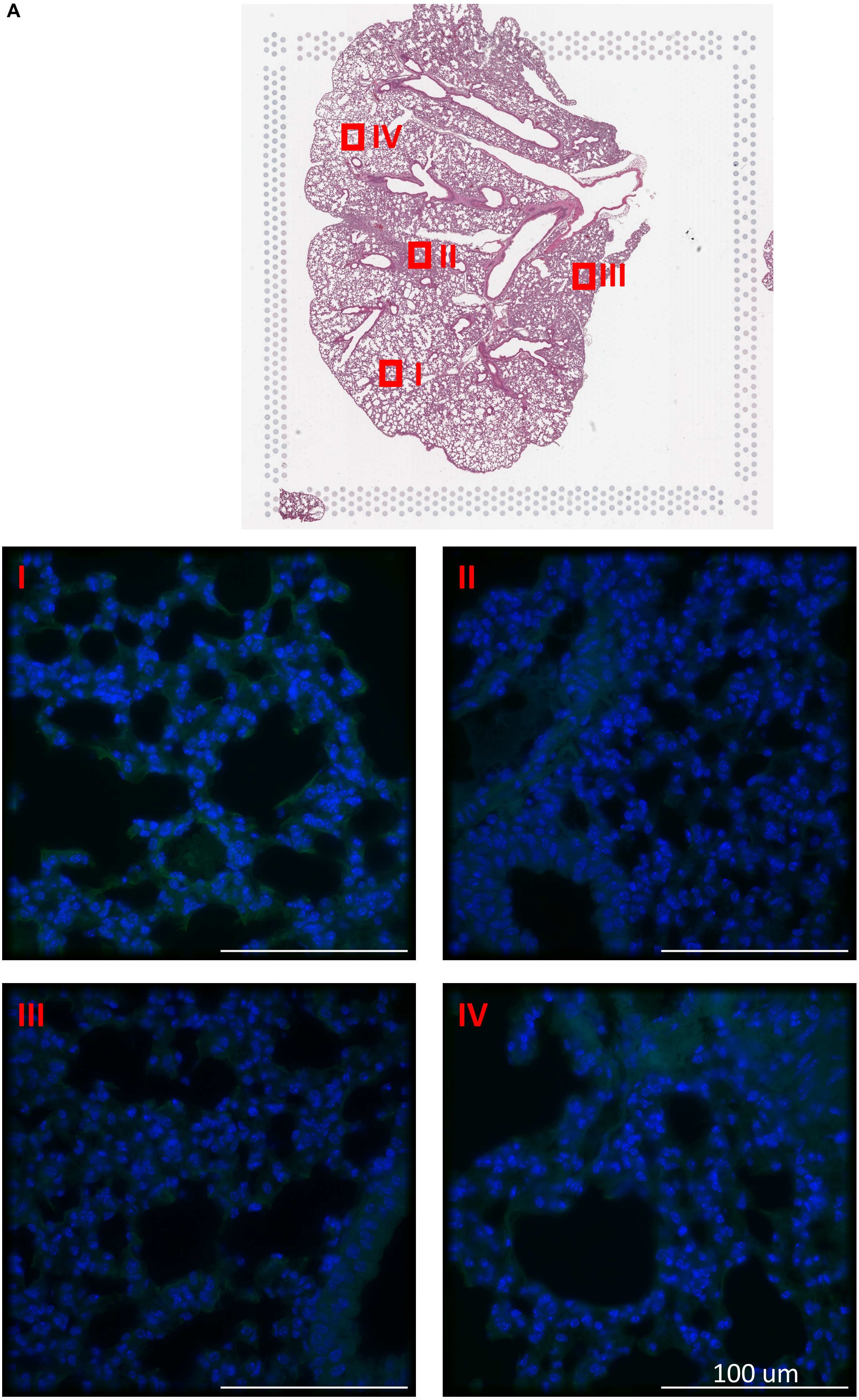


**Supplementary Figure 10**: **A**) Representative images of immunofluorescence staining in CTRL mice without infection (with specific antibody against *M. abscessus*, stained in green, and with Hoechst 33342, stained in blue) in four different areas (D IV) all captured in the subsequent slice of CTRL sample used for spatial transcriptomics (Scale bar, 100 μm).

**SUPPLEMENTARY FIGURE 11**

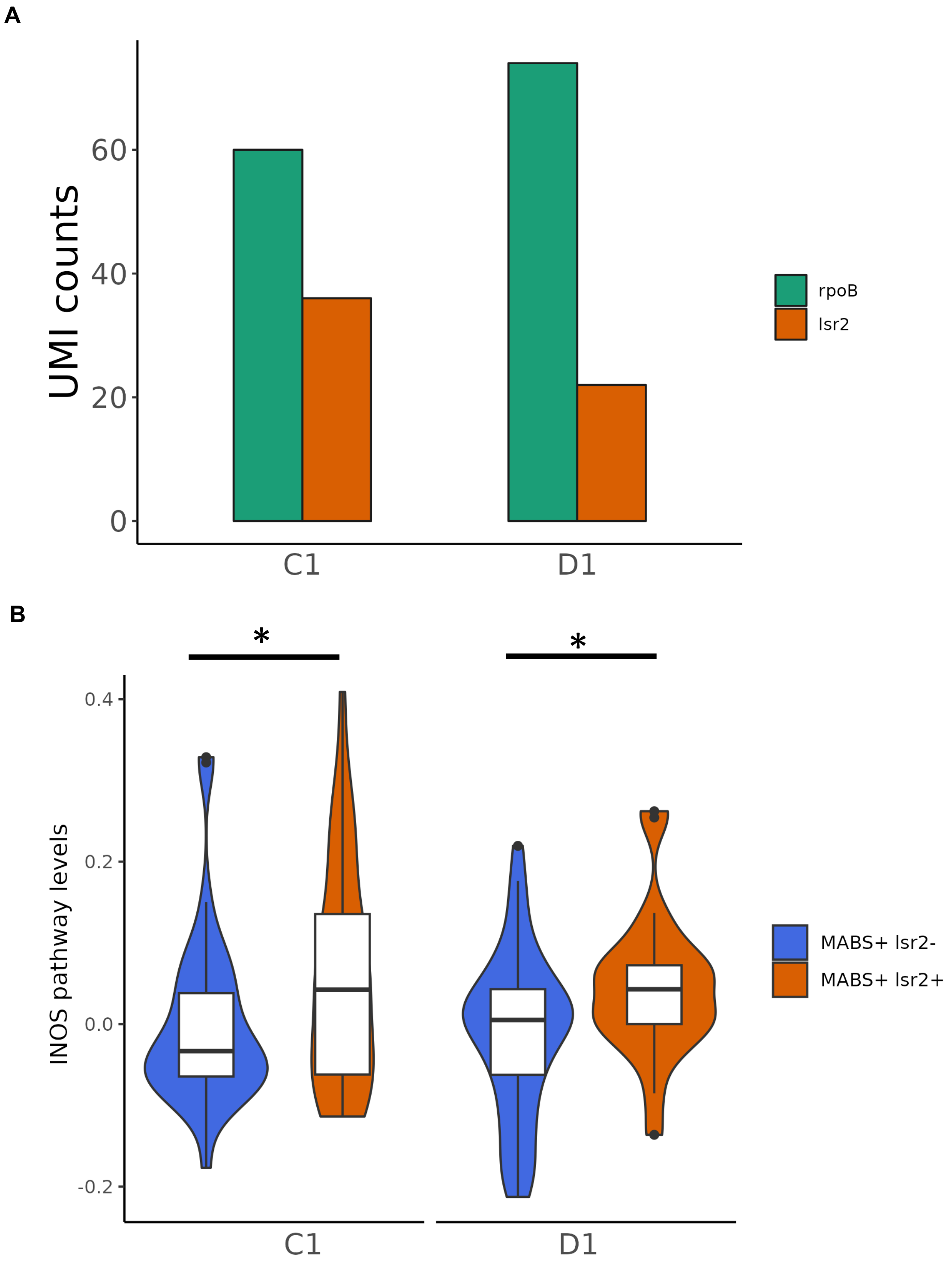


**Supplementary Figure 11: A**) Barplot showing the gathered UMI counts of rpoB and lsr2 probes in in C1 and D1 slices from murine lung with chronic infection. The y-axis shows the number of UMIs. The bars are color-coded by the respective probe. **B**) Violin plot for the levels of INOS pathway grouped according to MABS+lsr2- or MABS+lsr2+ Infection in C1 and D1 slices from murine lung with chronic infection. (Fisher's method for p-value aggregation , *p-value <0,05).
